# An approach to differentiating biological from chemical sulfide oxidation on corroded concrete sewer surfaces

**DOI:** 10.1101/2025.05.13.653906

**Authors:** Xin Huang, Yanpeng Yin, Zhiqiang Zuo, Tao Shi, Thomas Kuen, Zhiguo Yuan, Min Zheng

## Abstract

Hydrogen sulfide (H_2_S)-induced sewer corrosion largely impacts the longevity of infrastructure worldwide. The corrosion incorporates two pathways, i.e., chemical and biological oxidation of H_2_S to sulfuric acid; however, their rates are difficult to differentiate on corroded sewer pipelines, limiting understanding and application of effective strategies for corrosion control. Here, this study develops and reports a kinetic approach to determining rates of chemical and biological sulfide oxidation. By measuring the total sulfide uptake rate of sewer corrosion concretes, extensive experimental data was obtained for calibration and validation of the proposed kinetic model. The modelling results fit with measured data indicating that both chemical and biological sulfide oxidation can be well-described by the developed model. Model predictions revealed that along with sewer corrosion (i.e., pH decrease), chemical sulfide oxidation rate was significantly accelerated, along with an increase in biological rate. This result suggests that the chemical sulfide oxidation process should not be overlooked for seriously corroded concretes, and synergistic strategies to suppress both biological and chemical rates are required. Specifically, the chemical process can predominate sulfide oxidation with a gas-H_2_S concentration higher than 300 ppm. The work provides a useful tool for future corrosion studies and also valuable insights for sewer corrosion management practices.

**Synopsis:** This study reports a kinetic approach to determining chemical and biological H_2_S oxidation rates of sewer concretes.

**Graphic abstract:** 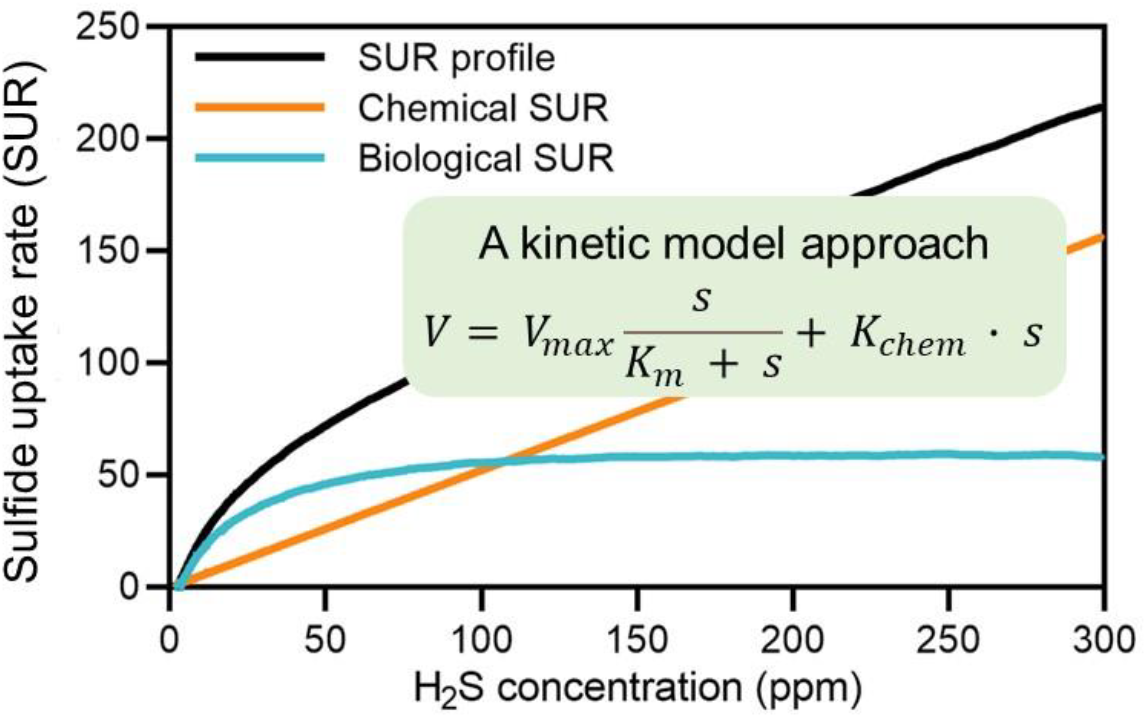

## 1 Introduction

Sewers are essential infrastructure systems, responsible for collecting and conveying sewage from various wastewater sources to treatment facilities, thereby safeguarding public health against waterborne diseases [1]. Investment in sewer systems is substantial; for instance, Brongers et al. estimated the value of sewer networks in the United States (US) at $1 trillion [2], and Jiang et al. reported that the total value of Australia’s sewers was $100 billion [3]. Nevertheless, sewer systems worldwide suffer from hydrogen sulfide (H_2_S)-induced corrosion, which incurs costs equivalent to 3.4% of the global GDP [4].

Sewer corrosion is primarily caused by H_2_S oxidation [5]. H_2_S can be generated by sulfate-reducing bacteria (SRB) in anaerobic biofilms and sediments [6], emitted from sewer sewage into the air phase, and undergo chemical and biological oxidation to form sulfuric acid through reaction with oxygen [7]. The accumulation of sulfuric acid on the surface of concrete pipes adversely affects pH, resulting in corrosion and ultimately pipe collapse [8, 9]. Initially, the pH of newly constructed concrete pipes is approximately 13. Sewer corrosion begins with the acidification of concrete pipe surfaces through chemical oxidation of H_2_S and carbonation of CO_2_ [10], which decreases pH to 9, thereby promoting the colonization of neutrophilic sulfide-oxidizing bacteria (SOB) [11]. Following that, both chemical and biological oxidation of H_2_S occur to form H_2_SO_4_ and reduce the surface pH to 4, allowing acidophilic SOB to dominate the corrosion process [12]. These acidophilic bacteria can thrive in highly acidic conditions, with pH levels ranging from 2 to 4 [13]. Studies have observed the presence of these acidophiles even at pH levels below 1 [9].

Identifying real-time corrosion rates is critical to prolong sewer service life. The corrosion rates can be reflected by measuring sulfide uptake rates (SUR), a quantitative, non-destructive and time-efficient method. SUR quantifies the H_2_S consumption rates on the surface of corroded concrete using a gas-tight reactor, establishing a direct correlation between H_2_S uptake rates and corrosion activity [14]. This technique relies on the principle that sewer corrosion results from the uptake and oxidation of sulfide to sulfuric acid (H_2_SO_4_). Due to the rapid adsorption of H_2_S and its direct conversion to H_2_SO_4_, this method can accurately quantify corrosion activity. Sun et al. determined the SUR of corroded sewer concrete before and after applying a bacteriostat, finding values of 83 and 13 mg-S m^−2^ h^−1^, respectively [14]. However, these existing SUR assessments primarily focused on corrosion activity resulting from the combined effects of chemical and biological H_2_S oxidation. Differentiation of biological from chemical sulfide oxidation should be essential for cost-effective control of sewer corrosion, yet no method is available to date. For instance, Li et al. manually elevated the H_2_S concentration to 1000 ± 100 ppm and observed a rapid onset of chemically induced corrosion [7].

Modelling is a critical tool in sewer management, providing valuable predictions for H_2_S production and oxidation. Various extensive models have been developed for this purpose. The Wastewater Aerobic/anaerobic Transformation in Sewers (WATS) model, for example, predicts H_2_S production in both anaerobic and aerobic processes [15]. Sharma et al. proposed a model to describe H_2_S production under dynamic sewer conditions [16]. The kinetics of biological H_2_S oxidation in sewer biofilms in the liquid phase were reported by Nielsen et al. [17], while the chemical H_2_S oxidation kinetics in aqueous solutions can be described by a first-order model [18]. These models were integrated into Sharma’s mathematical model [16]. Additionally, the comprehensive SeweX model has been successfully applied in the sewer industry to predict spatial variations in sulfide production and emission [19]. Recently, the Biofilm-Initiated Sewer process Model (BISM) was introduced to account for sulfide production along the rising main, incorporating the dynamics of biofilm activity [20]. Unfortunately, these previous models focus on addressing sulfur conversion in the liquid phase and do not consider the chemical and biological reactions occurring on corroded concrete surfaces.

This study aims to develop a tool to measure and estimate chemical and biological H_2_S oxidation rates on sewer concretes. A kinetic model was calibrated and validated using experimental data obtained from a series of corroded sewer concrete. By fitting data into the model, both the chemical and biological rates were quantified through calculated parameters. The derived results achieved high fidelities with the experimental data, indicating the acceleration of chemical and biological H_2_S oxidation rates as sewer corrosion processes. Subsequently, the chemical and biological contributions to the total rate were predicted for corroded concretes under different corrosion stages, identifying the principal H_2_S oxidation pathway. The findings suggest that more attention should be paid to the predominant pathway for sewer corrosion control.

## 2 Model method description

According to the SUR method, gas-H_2_S concentration decreases along with time (Figure 1A). With H_2_S data monitoring, the total rate for sulfide uptake can be determined (i.e., SUR profile in Figure 1B). However, the total rate includes both rates of chemical and biological processes (i.e., chemical and biological SUR conceptually in Figure 1B), though they cannot be directly differentiated. Here, an integrated sulfide oxidation model describes both chemical and biological processes, as depicted in Eq. (1).

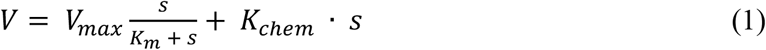

where *V* is measured SUR profile of sewer concrete (mg-S·m^−2^·h^−1^), *V*_max_ is the maximum specific growth rate of SOB on the concrete (mg-S·m^−2^·h^−1^), *K*_m_ is the substrate half-saturation constant (ppm), *s* is the H_2_S concentration (ppm), *K*_chem_ is the chemical oxidation rate constant (mg-S·m^−2^ h^−1^ ppm^−1^).

**Figure 1.**
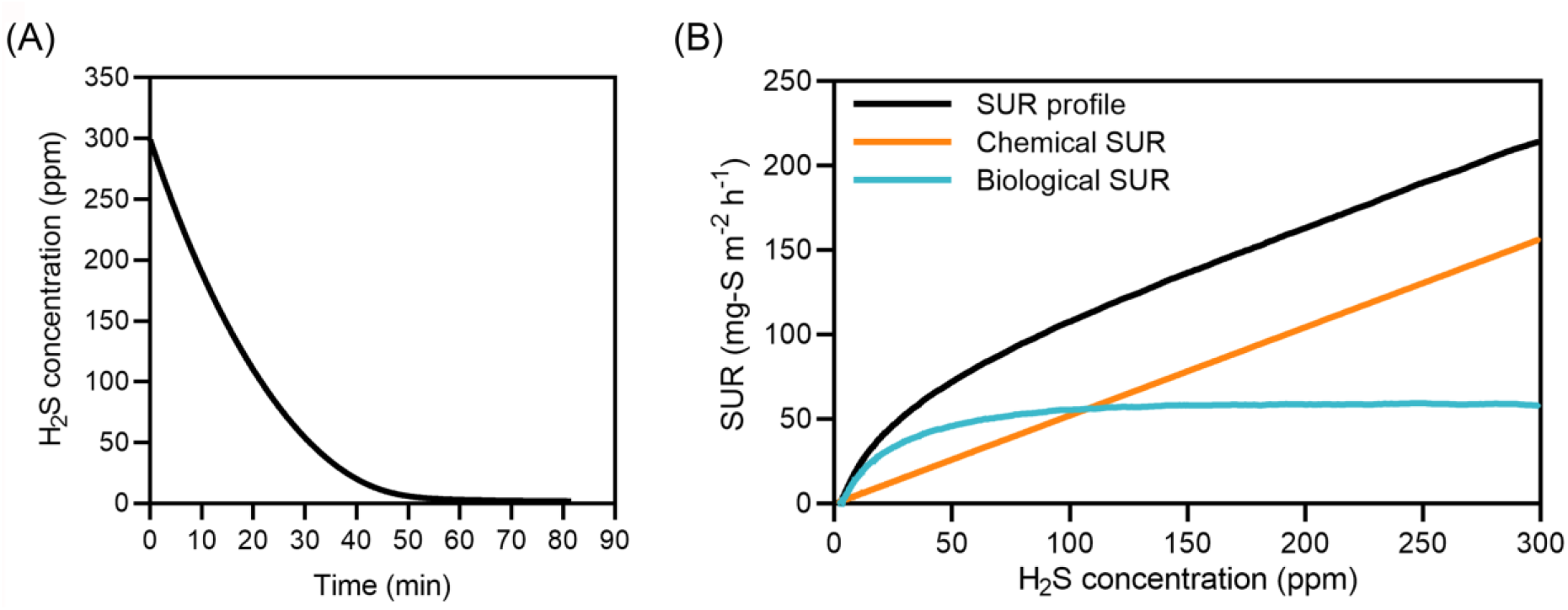
(A) The typical profile of gas-H_2_S concentration decrease due to sulfide uptake in the short-term test. (B) Profile of total sulfide uptake rate (the sum of biological and chemical rates) based on measured gas-H_2_S concentration versus time, and the hypothetical profiles of biological and chemical rates.

With experimental verification below (see section 4.1), the first-order kinetic equation was used to describe the chemical sulfide oxidation process. Since biological sulfide oxidation is driven by microbial activity of SOB that use H_2_S as a nutrient source, the Monod equation, widely used to describe kinetics in many biological treatment processes [17], was proposed to describe the biological sulfide oxidation process.

## 3 Experimental designs

### 3.1 Source of corroded concrete coupon

Concrete coupons of various ages, along with a virgin coupon (VC), were utilized for SUR measurements in the present study in order to prove the developed model. Each concrete coupon was made by casting two concrete blocks into a stainless-steel frame and fixing them with epoxy [14]. 2 vinyl tubes (Cole Parmer, 3/8 ID, 1/2 OD, 1/16 Wall)) were installed at the bottom of each stainless-steel frame. The tubes allowed for interconnection between the coupons. The upper surfaces of the concrete coupons were exposed to sewer air, leading to H_2_S-induced deterioration. All concrete blocks were fabricated in accordance with previous protocols [21] and complied with the Australian standard [22], with dimensions of 100 mm (length) × 70 mm (width) × 50 mm (thickness). The mechanical properties, including compressive strength, hardened density, drying shrinkage, and permeable voids, of the manufactured concrete were consistent with those of real sewer pipes [23]. To expedite the neutralization of fresh concrete by H_2_S and CO_2_, a 10 wt% H_2_SO_4_ solution was applied to the surface of the coupons, as previously described [24].

The treated coupons were subsequently placed in a corrosion chamber to undergo the sewer corrosion process. The corrosion chamber was constructed to simulate a typical sewer corrosion environment for the manufactured concrete coupons (Figure. S1). The corrosion chamber had dimensions of 130 cm (length) × 120 cm (width) × 100 cm (thickness). The corrosion chamber housed a continuous circulation system comprising two 5 L wastewater tanks and eight drawers (56 cm × 55 cm × 9 cm) driven by two peristaltic pumps (BT-600EB, Chongqing Jieheng Peristaltic Pumps Co., Ltd) operating at 100 rpm, with a total volume of 50 L. The flowing water maintained the chamber’s relative humidity (RH) at approximately 99%.

A 50% HCl solution and a 70 g/L Na_2_S solution were mixed in a 10 L carboy within the chamber using two peristaltic pumps (SEKO, PR-4 detergent pump) to produce H_2_S gas. Two H_2_S loggers (IPX H_2_S PPM, Acrulog™) were used to monitor the chamber’s H_2_S concentration, which was maintained at 50 ppm. The recommended H_2_S concentration was provided by our industry partners. The pH of the wastewater was monitored using pH probes (InPro3250i/SG/225, Mettler Toledo) installed in the wastewater tanks. When the pH dropped below 7, NaOH solution (40 g/L) was introduced into the wastewater tanks. The chamber’s air temperature was maintained at 23 °C by centralized air conditioning in the laboratory. Prior to installation on the racks in the chamber, each neutralized coupon received an even spray of 10 mL of wastewater to facilitate microbial inoculation. The exposed concrete surfaces faced downward on the racks to simulate the crown region of sewer pipes, a known corrosion hotspot in real-world scenarios [25]. Vinyl tubes connected the concrete coupons inside the chamber, enabling the flow of cooling water between them. This arrangement maintained the coupons’ temperature slightly below that of the chamber air, leading to the formation of condensation on the coupon surfaces. Coupons were incubated in the corrosion chamber for varying durations before undergoing SUR tests to replicate sewer pipes with different service ages.

### 3.2 Characterization of corroded concrete coupon

Three corroded coupons were retrieved from the corrosion chamber for a series of characterization tests to assess their corrosion status prior to conducting SUR tests. Surface pH measurements, surface area scanning, and visual observation were conducted after retrieving the corroded coupons from the corrosion chamber. Prior to pH measurement, 1 mL of milli-Q water was applied to the coupon surface, following the procedure outlined in a previous study [3]. The surface pH of the coupons was measured using a flat surface pH probe (RapidRH® portable pH kit, Wagner) after a 1-minute interval. pH values from three different locations on each coupon surface were recorded to calculate the average surface pH of the coupon. Following pH measurement, a 3D scanner (EinScan-SE, Shining 3D) was used to analyze the surfaces of the coupons. The exposed concrete surfaces were scanned multiple times and from various angles to generate high-quality digital images of each coupon using the software (EXScan S). Subsequently, the surface area of the corroded concrete was calculated based on the digital images using the software’s built-in model. Table 1 presents the properties of the tested coupons.

**Table 1.**
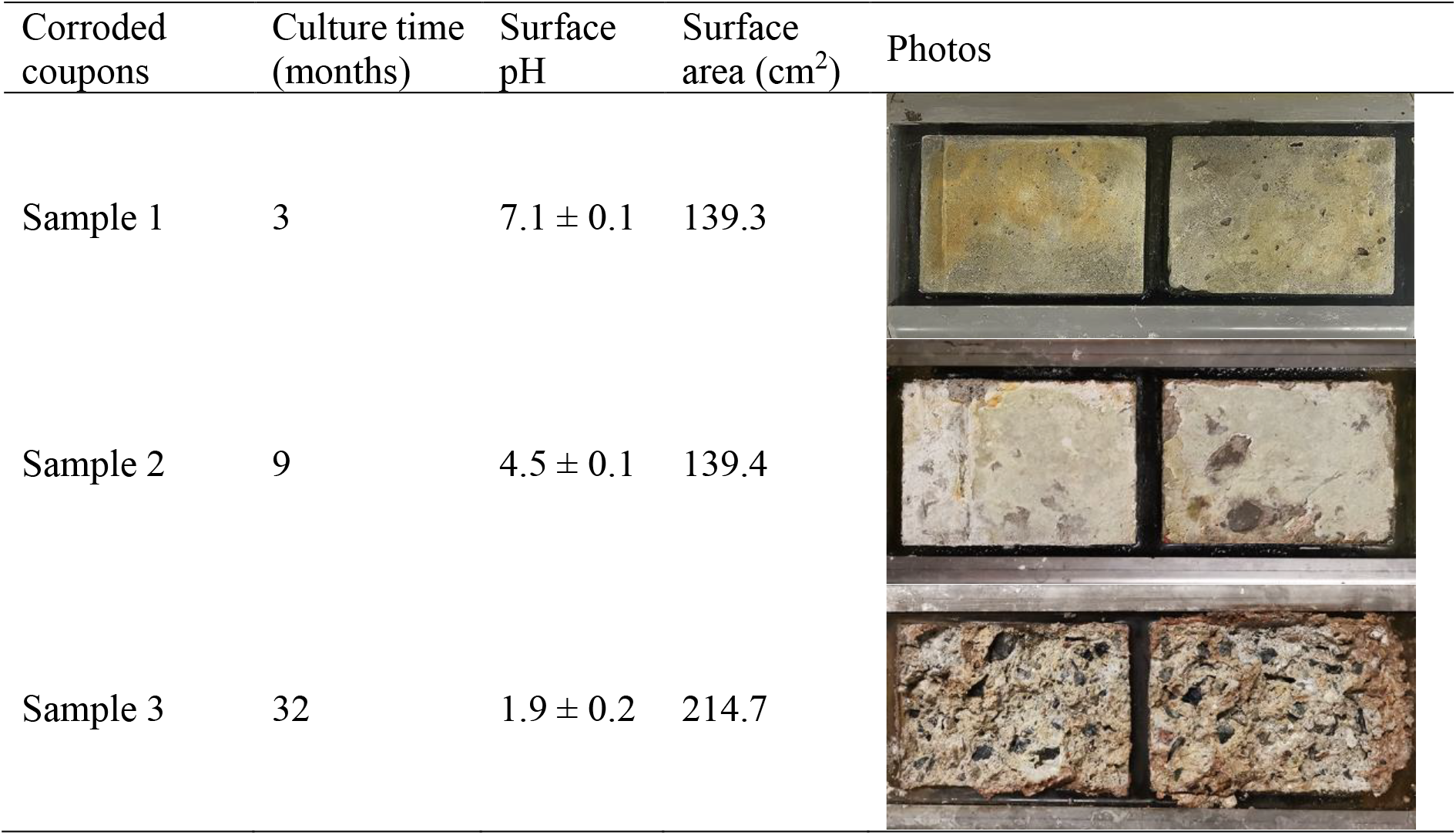
Characteristics of the corroded coupons.

| Corroded coupons | Culture time (months) | Surface pH | Surface area (cm<sup>2</sup>) | Photos |
| --- | --- | --- | --- | --- |
| Sample 1 | 3 | 7.1 ± 0.1 | 139.3 |  |
| Sample 2 | 9 | 4.5 ± 0.1 | 139.4 |  |
| Sample 3 | 32 | 1.9 ± 0.2 | 214.7 |  |

A virgin coupon (VC), which was neutralized but not incubated in the chamber afterwards, was also included in this study as the non-corroded control coupon. The characterization tests were also conducted to assess the properties of VC. The surface pH of the VC was adjusted to 7.7, and its surface area was comparable to that of sample 1.

### 3.3 Sulfide uptake rate (SUR) experiments

SUR profiles of the tested coupons were recorded following the methodology outlined in a previous study, with modifications made to validate the proposed kinetic model [14]. After collection from the corrosion chamber, each coupon was placed onto a gas-tight glass reactor (30 cm × 12.5 cm × 11 cm). Before conducting the SUR tests, the RH of the SUR reactor was raised to 100% by spraying 3 mL of Milli-Q water into the reactor. H_2_S was generated in a 100 mL bottle by mixing a 22 g/L Na_2_S solution with HCl solution. Subsequently, the generated H_2_S was gradually transferred to the SUR reactor via a silicone tube. Two small fans (YX 2503, Sirocco) inside the reactor were utilized to ensure uniform air distribution within the reactor. H_2_S sensors (App-Tek OdaLog® Logger L2) were installed on the reactor to monitor the decrease in H_2_S levels from 300 ppm to 0 ppm. The sampling time interval of the H_2_S sensors was set to 5 seconds. Each coupon was measured three times to ensure the reproducibility of the results. The calculated slopes of H_2_S profiles indicated the H_2_S uptake rates. The surface-specific H_2_S uptake rate was calculated by the slope and the surface area of the exposed concrete.

The overall design of all SUR tests is summarized in Table 2. Batch test I was conducted to validate the first-order chemical sulfide oxidation equation. VC and an acrylic board were used to record chemical SUR under various pH and RH conditions. In the third test in this set, the surface pH of VC was adjusted from 7.7 to 5.0 by adding a 10 wt% H_2_SO_4_ solution to the coupon, and the RH in the SUR reactor was reduced from 100% to 50% using a saturated Mg(NO_3_)_2_ solution. For the acrylic board, the test was performed at a surface pH of 7.0 and an RH of 100%. Batch test II was carried out on corroded coupons (no. 1, 2, and 3) at 100% RH, and the obtained data was used to validate the integrated model, which includes both chemical and biological processes. Additionally, we also exploited data from a previous pilot study to validate the proposed model in this work [26].

**Table 2.** Batch tests design of SUR measurements on solid surfaces.

| No. | Tested materials | Surface pH | RH in the reactor | Objectives |
| --- | --- | --- | --- | --- |
| **Batch test I** | Virgin coupon | 7.7 ± 0.1 | 100% | Validation of the chemical oxidation equation |
|  | Virgin coupon | 7.7 ± 0.1 | 50% |  |
|  | Virgin coupon | 5.0 ± 0.1 | 100% |  |
|  | Acrylic board | 7.0 ± 0.1 | 100% |  |
| **Batch test II** | Corroded coupon 1 | 7.1 ± 0.1 | 100% | Validation of the combined chemical and biological model |
|  | Corroded coupon 2 | 4.5 ± 0.1 | 100% |  |
|  | Corroded coupon 3 | 1.9 ± 0.2 | 100% |  |

### 3.4 Data analysis

The measured data were processed with Excel (2016, Microsoft) to calculate surface-specific SUR. The SUR profiles were fitted to our model using MATLAB (R2023a). Simple linear regression was applied to the SUR profiles of VC and the acrylic board, while nonlinear least squares fitting was employed for SUR profiles of corroded coupons. The 95% confidential ellipses were calculated and plotted using MATLAB. Other figures were generated using Prism (9.3.1, GraphPad Software, LLC) and Origin. The standard deviation of triplicate results was calculated using a built-in function of Prism.

## 4 Results

### 4.1 SUR test I: calibration of chemical oxidation kinetics

The chemical oxidation process of H_2_S was investigated using the VC group, which was neither inoculated with wastewater nor cultured in the corrosion chamber, ensuring no SOB present on its surface. Consequently, the measured SUR only reflects the sole chemical oxidation of H2S. As shown in Figure 2, triplicate SUR measurements were conducted under various conditions. Under 100% RH, the SUR profiles of VC exhibited a proportional relationship with H_2_S concentrations (Figure 2: A1-A3). Simple linear regression analysis of the SUR data indicated a good fit, suggesting that the chemical oxidation of H2S on the coupon surface follows first-order kinetics. Subsequent SUR measurements were also conducted under varying RH and pH conditions to simulate chemical H_2_S uptake in diverse sewer systems (Figure 2B-C). This is essential due to the variations in RH and surface pH among sewer pipes in different regions [27]. SUR measurements were further performed on an acrylic board, and the results confirmed the same chemical H_2_S oxidation profiles of solid materials (Figure 2D). All SUR profiles showed good fitting profiles through linear regression analysis. Therefore, it can be concluded that chemical oxidation of H2S on solid surfaces follows the first-order kinetics.

**Figure 2.**
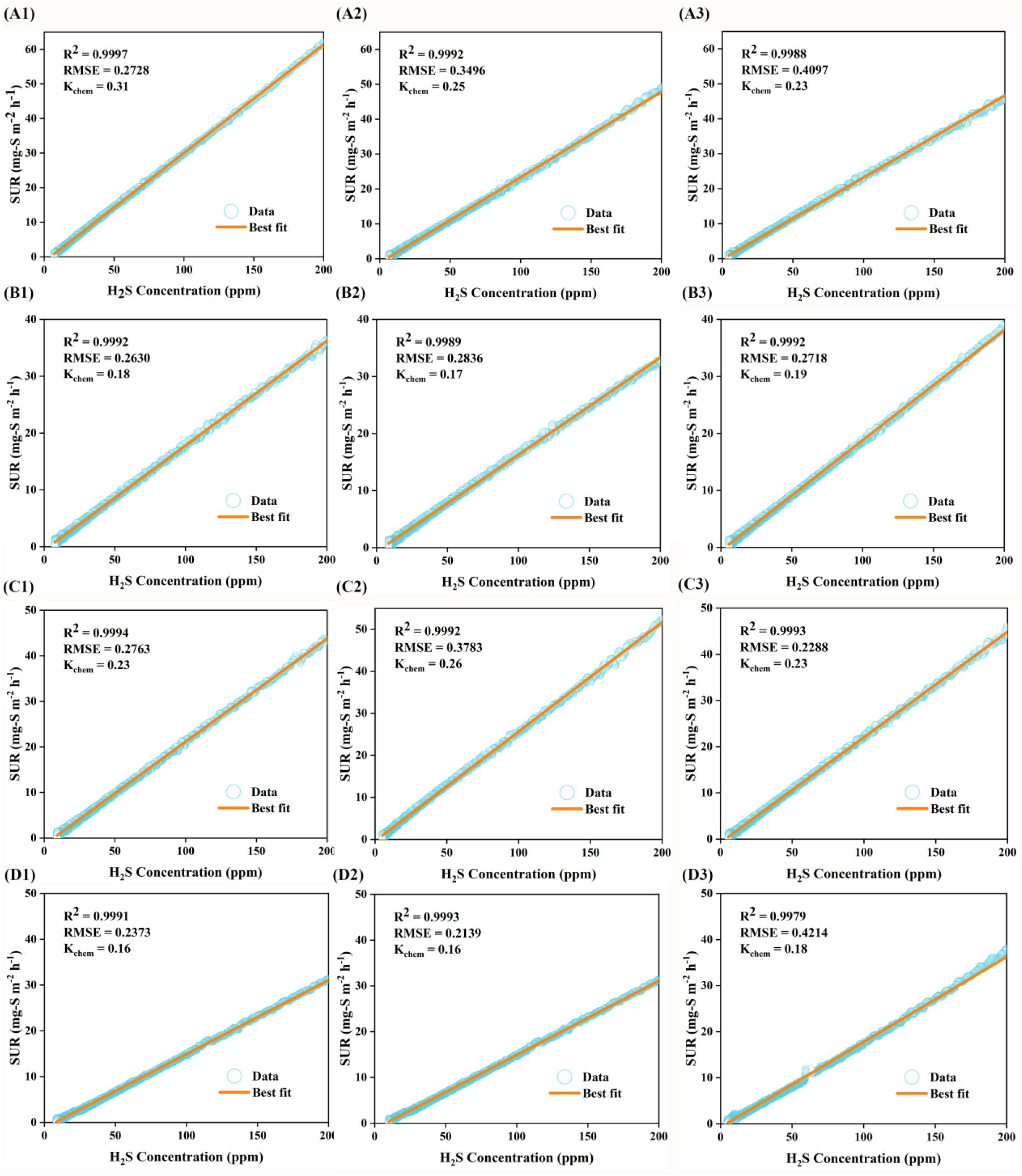
Chemical SUR profiles of solid surfaces and their simple linear fitting results. (A1-3) are the VC value of the surface pH of 7.7 in 100% RH atmosphere. (B1-3) are the measurement of VC with a surface pH of 7.7 in 50% RH atmosphere. (C1-3) are the results of VC with a surface pH of 5 in 100% RH atmosphere. (D1~3) are measurements on an acrylic board with a surface pH of 7 in 100% RH atmosphere.

### 4.2 SUR test II: validation of the integrated chemical and biological kinetics

The total SUR for corroded coupon samples was measured at H_2_S concentrations ranging from 0 to 200 ppm in 100 % RH air. The experimental results are displayed in Figures 3A, 3B and 3C. The total SUR rapidly increased at an H_2_S concentration of less than 20 ppm, then gradually rose with an H_2_S concentration of around 100 ppm. At a higher H_2_S level (> 100 ppm), the total SUR profiles obtained in Figure 3 can be described by the simple linear regression. The results indicated that the increase in total SUR was mainly due to the enhancement of chemical SUR because the biological rate could be maximized in the Monod equation with high substrate H_2_S concentrations. The total SUR profiles of samples 1, 2, and 3 differed from that of VC when H_2_S levels were below 100 ppm, which should be mainly attributed to biological H_2_S oxidation. Among these three samples, the samples with lower pH values were associated with higher total SUR at the same H_2_S level. For instance, coupons corroded for 3 and 9 months (samples 1 and 2, pH at 7.1 and 4.5, respectively) showed the total SUR of 38 mg-S·m^−2^·h^−1^ and 151 mg-S·m^−2^·h^−1^ at 200 ppm (Figures 3A and 3B), respectively. After 32 months of corrosion (sample 3, pH at 1.9), the total SUR markedly increased to 366 mg-S·m^−2^·h^−1^ (Figure 3C).

**Figure 3.**
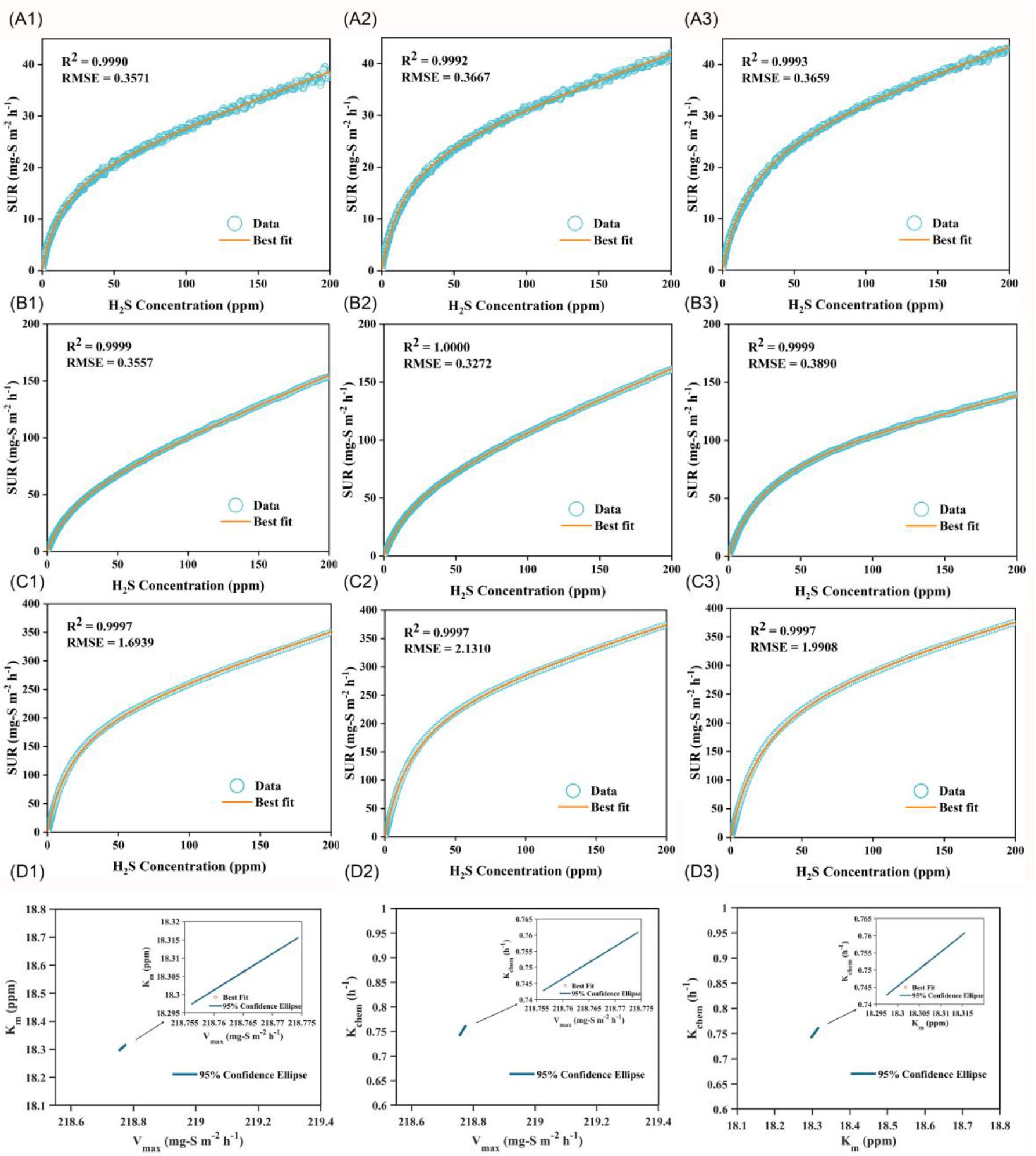
Total SUR and fitting curves of the corroded coupons. (A), (B) and (C) are the triplicate results of samples 1, 2, and 3. (D) shows the 95% confidence ellipses of the calculated parameters from (C1) as an example.

Furthermore, data fitting was performed on the experimental results by using the developed model. Overall, the model well described all SUR measurements under these different conditions, thereby effectively validating the model. To avoid over-data-fitting, correlations among critical model parameters (i.e., between *K*_m_ and *V*_max_, *K*_chem_ and *V*_max_, and *K*_chem_ and *K*_m_) were examined with 95% confidence ellipses (Figure 3D). Taking the measurement for sample 3 as an example (e.g., Figure 3 C1), though the narrowness of ellipses correlated with the parameters, the range was remarkably narrow, indicating the effectiveness of the calibrated parameters.

### 4.3 Values of calculated kinetic parameters

Figure 4 shows the values of calibrated parameters *K*_chem_, *V*_max_, and *K*_m_ for the three corroded coupon samples. The *K*_chem_ values of samples 1, 2, and 3 were 0.09 (± 0.003), 0.41 (± 0.112), and 0.64 (± 0.052) mg-S·m^−2^ h^−1^ ppm^−1^, respectively. The *K*_chem_ significantly increased with the progression of sewer corrosion with the sample pH decreasing from 7.1 to 4.5 and 1.8 (Figure 4A). This result suggested that heavily corroded sewer pipes might experience intense chemical H_2_S oxidation. The biological rate also considerably increased as the corrosion stage progressed along with the pH decrease. The *V*_max_ value rose from 24 to 262 mg-S·m^−2^·h^−1^ from samples 1 to 3 (Figure 4B), underscoring a substantial role of microbial activity. Furthermore, the *K*_m_ values for the three samples were estimated to be about 20 ppm showing a similar mass transfer rate of H_2_S to biological oxidation.

**Figure 4.**
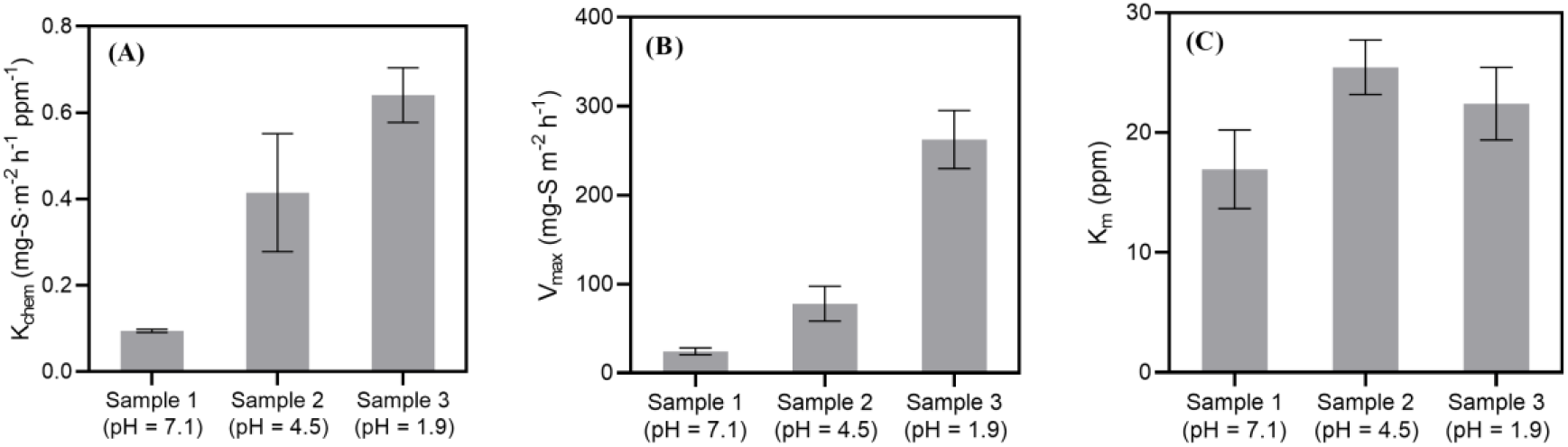
The calculated parameters of Eq. 1 from the total SUR of samples 1, 2, and 3. (A) compares the chemical H_2_S oxidation constants. (B) shows the calculated maximum biological SUR, and the substrate half-saturation constants are plotted in (C).

### 4.4 Contribution of chemical and biological sulfide oxidation under different H_2_S concentrations

The developed model and calibrated parameters were used to quantitatively assess rates of chemical and biological H_2_S oxidation under various H_2_S concentrations of 15, 50, 100 and 300 ppm, which encompass a wide concentration range in real sewers [7, 28]. The proportion of biological rate in the total sulfide oxidation rate decreased with the increased H_2_S concentration. At a low H_2_S level (< 100 ppm), the proportion of biological oxidation rate in the total rate exceeded that of chemical reaction (Figure 5). The rate of biological oxidation can predominate sulfide oxidation when H_2_S is relatively low and generally reach the maximal level at H_2_S of ~100 ppm. After that, chemical and biological oxidation had nearly equal proportions in the total sulfide oxidation at H_2_S concentrations of about 300 ppm. This result underscores the overlooked high contribution of chemical oxidation to sewer corrosion at high H_2_S concentrations.

**Figure 5.**
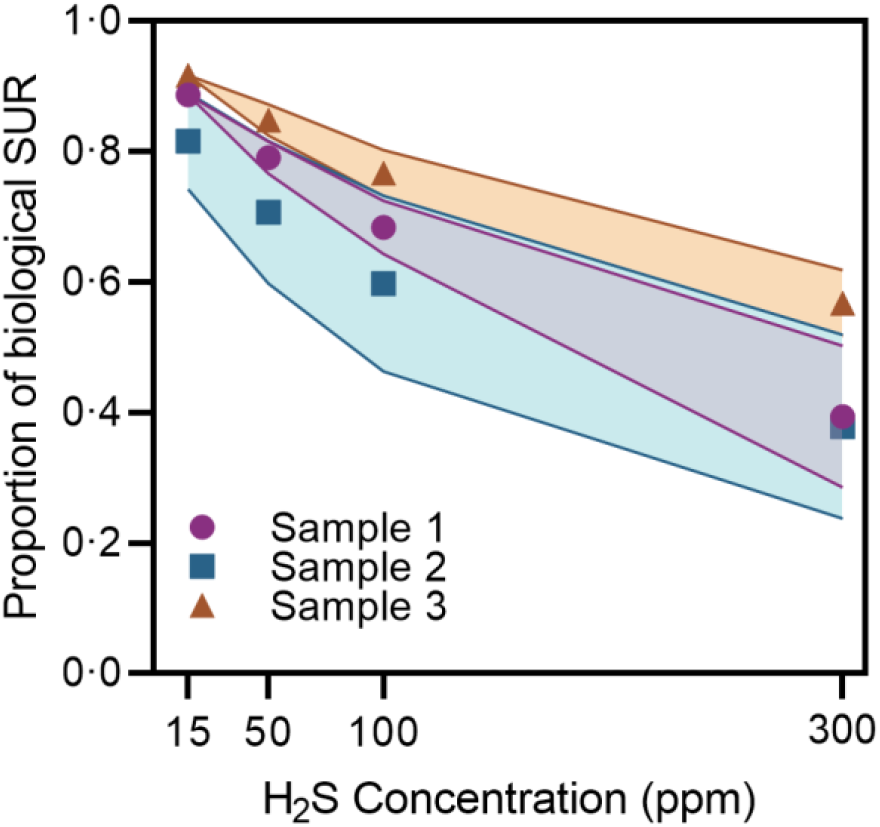
The simulated proportion of biological rate in the total sulfide oxidation rate at H_2_S levels of up to 300 ppm. The boundary of the shaded area indicates the error bar of the corresponding samples.

## 5 Discussion

### 5.1 A new approach for determining chemical and biological H_2_S oxidation

The SUR has long been recognized as an effective indicator of corrosion activity. However, it failed to differentiate between chemical and biological oxidation pathways in previous studies [14, 29]. This study proposes a new kinetic model approach, validated through extensive experiments with model fitting, to more accurately estimate chemical and biological rates. The model was further validated using H_2_S consumption data from a pilot sewer pipe study [26], which demonstrated good fits with data at H_2_S concentrations up to 900 ppm (Figure S2).

This study determined a first-order kinetic equation for chemical H_2_S oxidation on sewer concrete surfaces (Figure 2), a finding that contrasts with previous studies reporting that chemical sulfide oxidation in aqueous phases followed the n^th^ order kinetics (n = 0.90-1.38) [30, 31]. The difference is likely due to higher sulfide concentration (up to 8 g S m^−3^) in solution affecting H_2_S mass transfer than on solid surface. Indeed, by examining chemical H_2_S oxidation on concrete surfaces across a wide H_2_S concentration range of 10 ppm to 200 ppm and 500-1500 ppm [7], a recent study reported the first-order kinetic pattern between H_2_S of 10 ppm and 125 ppm, followed by exponentially increased rate at a higher H_2_S concentration. The kinetic discrepancy for chemical sulfide oxidation needs to be carefully considered in future corrosion studies, particularly at extremely high H_2_S concentrations.

### 5.2 Accelerated corrosion by both chemical and biological sulfide oxidation

The corrosion process can become worse along with the progression of corrosion, as increasing rate of biological H_2_S oxidation often occurs along with SOB proliferation versus time, namely acceleration of sewer corrosion [7, 15]. As estimated in this work, the maximal biological H_2_S oxidation rate significantly increased in the corroded samples with low pH (Figure 4B). The evolution of microbial communities and accumulation of biomass on coupon surfaces were believed to play a crucial role [1]. For instance, neutrophilic SOB tend to dominate with pH decrease down to 4, while acidophilic SOB could proliferate at lower pH. Analysis of microbial compositions in samples 2 and 3 through 16s rRNA gene amplicon sequencing revealed that the relative abundance of acidophilic SOB *Acidithiobacillus* in the total community increased from 56% to 67% with the surface pH decreasing from 4.5 to 1.9 (Figure S4). Major microbes present at pH 4.5, like *Metallibacterium* (18%) and *Dyella* (6%), were eradicated at pH 1.9. At the extremely low pH, iron-oxidizing archaea *Ferroplasmaceae* showed a very high abundance of 32% in the total community, while its role remains unclear [32].

Interestingly, chemical sulfide oxidation also had a significantly higher rate in the corroded samples with lower pH, indicating that the enhanced chemical reaction could occur during accelerated sewer corrosion. One possible reason is that H_2_SO_4_ produced on concrete surfaces can leach metals, and metal ions in corrosion layer can catalyze chemical sulfide oxidation [33–36]. Another reason could be that corrosion induces roughness and porosity in concrete, augmenting its chemical adsorption capacity to H_2_S [37]. In our further SUR test, the *K*_chem_ value of a sterilized corroded coupon sample was low compared to a non-sterilized coupon (Figure S3). The disparity may suggest different H_2_S adsorption capacities between sterilized and non-sterilized coupons.

### 5.3 Implications for corrosion management

Corrosion control is essential for effective sewer management, and various strategies have been developed to address this issue. Some strategies aim to decrease H_2_S concentration in the sewer air, e.g., injection of air or oxygen [29, 38] and dosing of chemicals such as nitrate [39], iron salts [40], alkali [41]. Others target microbial actions, e.g., suppressing SRB activity with free nitrous acid [42], free ammonia [43, 44] and controlling SOB growth by surface washing [21], biocidal spraying [45], etc. Considering the developed model, which reveals varied chemical and biological rates in different corroded pipelines, application of those strategies could be more effectively tailored. As known, sulfide-induced corrosion development involves early-middle-late stages with continuous pH decrease. In the early corrosion scenario, it is not recommended to use SOB-controlled strategies because of the minor contribution of biological sulfide oxidation. Instead, reducing H_2_S accumulation in liquid phase is more effective, as chemical oxidation mainly relies on H_2_S concentration. However, in the mid or late corrosion scenarios, both biological and chemical sulfide oxidation play crucial roles. Given that the H_2_S half-saturation constant for biological sulfide oxidation on corroded concretes is relatively low (~20 ppm), H_2_S concentration must be decreased at a level lower than H_2_S half-saturation constant. This is challenging because the current strategies cannot completely eliminant H_2_S concentration. In real sewer environment, sulfide concentration at several ppm-levels in wastewater has the potential to release several hundred ppm of gaseous H_2_S [46]. Thus, directly suppressing SOB growth through surface washing or biocidal spraying may be more direct and effective.

## 6 Conclusions

This study reports an integrated modelling approach to determining chemical and biological sulfide oxidation rates of corroded concrete samples. Conclusions include:

- Chemical sulfide oxidation on solid surfaces can be described by first-order kinetics. Combined with the Monod equation for the description of biological oxidation, the integrated model can enable modelling of the whole sulfide uptake process.
- Substrate H_2_S half-saturation constant for biological sulfide oxidation on corroded concretes was estimated to be 16.9 – 25.4 ppm.
- Biological and chemical sulfide oxidations have higher rates in more heavily corroded samples, suggesting both processes may proceed with the acceleration of corrosion in real sewer networks.
- Biological oxidation predominates in sulfide oxidation at low H_2_S concentrations (e.g., < 100 ppm), whereas chemical oxidation becomes more significant at higher H_2_S levels (e.g., >300 ppm).

## Acknowledgements

This work was financially supported by an Australian Research Council (ARC) Linkage Project (LP190101262). Xin Huang acknowledges the Chinese Scholarship Council for supplying the Living Allowance Scholarship. Dr. Zhiqiang Zuo acknowledges the support of the International Postdoctoral Exchange Fellowship Program (China). Prof Zhiguo Yuan is Global STEM Scholar funded by the Innovation and Technology Commission of the Government of the Hong Kong Special Administrative Region.

