## Supplementary material for "An approach to differentiating biological from chemical sulfide oxidation on corroded concrete sewer surfaces": Supplimental Information

# 6 SI

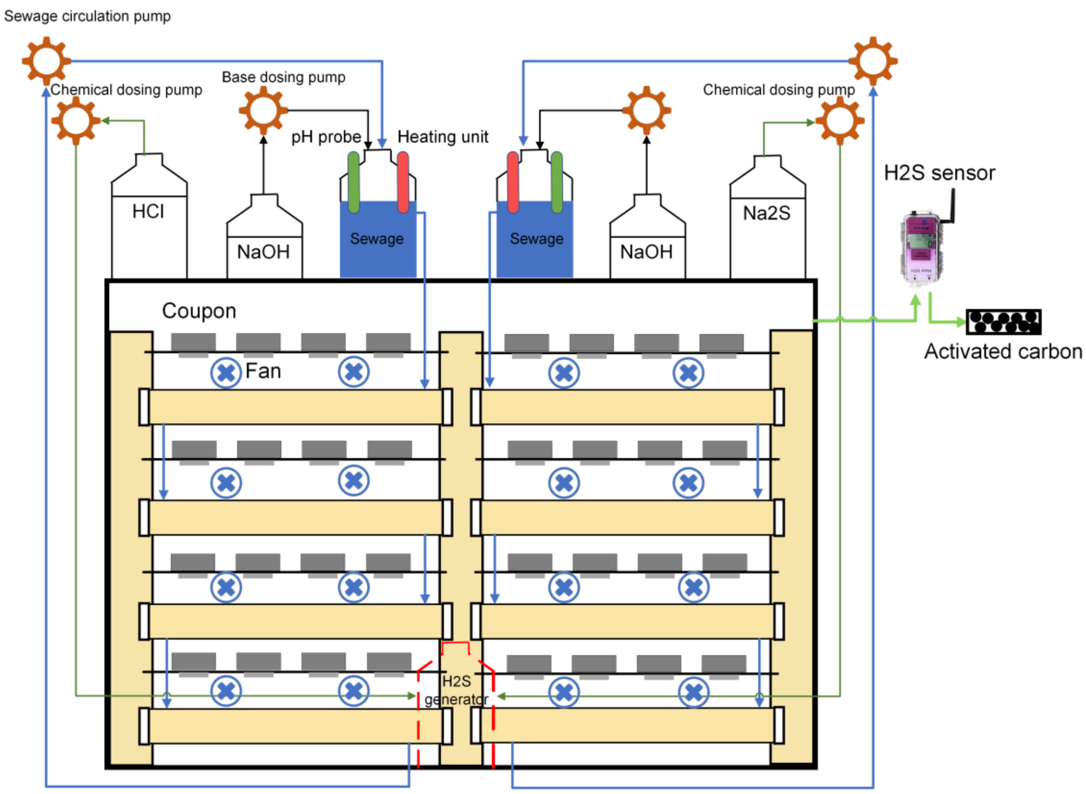


**Figure S1.** The design of a sewer corrosion chamber.

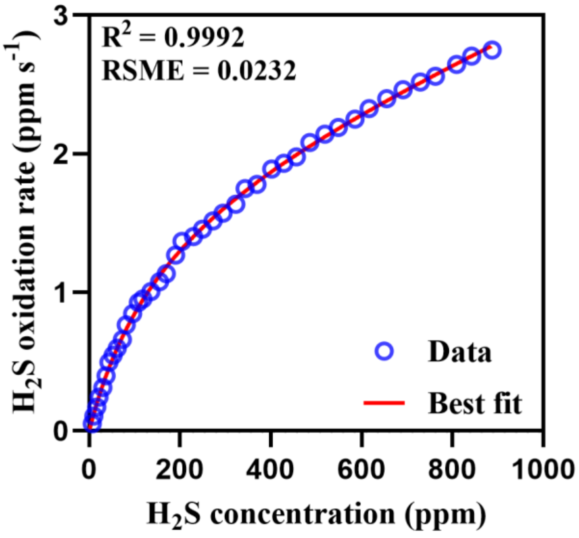


**Figure S2.** The total SUR of a pilot sewer pipe (data adapted from Vollertsen et al., 2008).

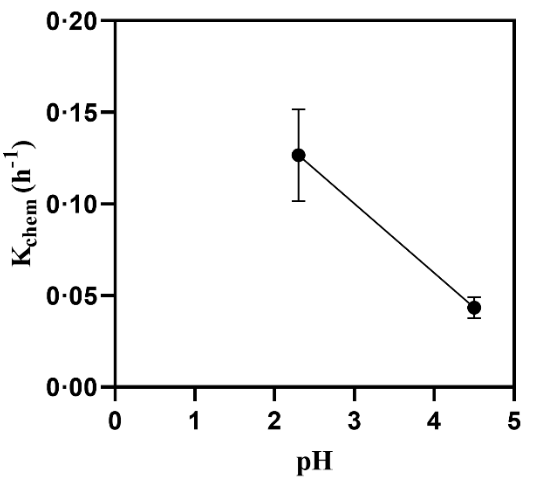


**Figure S3.** Chemical SUR of sterilized corrosion layers. The SUR tests were performed on the extracted corrosion layers from samples 2 and 3 laid onto a plastic board. The corrosion layers were sterilized in a 120℃ autoclave for 30 min before being covered onto the board.

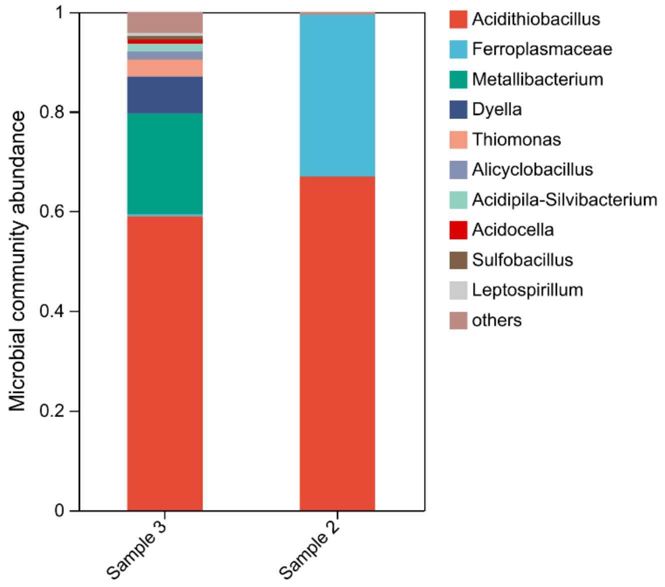


**Figure S4.** The microbial communities of samples 2 and 3.

Following the SUR measurements, samples 2 and 3 were destructively analyzed to obtain the corrosion layers and extract microbes. The microbial communities of the corroded coupons were analyzed using 16S rRNA analysis. Due to the absence of inoculated microbes on VC and the lack of extractable corrosion layers on sample 1, microbial community analysis was conducted solely for samples 2 and 3. Before DNA extraction, cells must be extracted from the corrosion layers as per the methodology outlined in previous research (Jiang et al., 2016). In brief, cells were liberated from the corrosion layer via ultrasonic treatment and collected in a sucrose solution. Subsequently, a 0.8% w/v NaCl solution was employed to wash the cells. The subsequent DNA extraction procedure utilized the FastDNA SPIN Kit for Soil (MP

Biomedicals, USA) according to the manufacturer's instruction, and the quality of the obtained DNA was examined by a NanoDrop microvolume spectrophotometer (ND-1000, Thermo Fisher, USA).

Extracted DNA samples were analysed by Shanghai Majorbio Bio-pharm Technology Co., Ltd (Shanghai, China) with 16S rRNA gene amplicon sequencing (Illumina). PCR amplifications were conducted in ABI GeneAmp&reg 9700 using TransStart Fastpfu DNA Polymerase (TransGen AP221-02). To detect the microbes from corroded sewers, modified universal primers 926F (50- AAACTYAAAKGAATTGRCGG -30) and 1392R (50-ACGGGCGGTGWGTRC -30) were used to target the V6-8 regions of the 16S rRNA gene. Sequences were processed with Quantitative Insights Microbial Ecology II (QIIME II) to remove low-quality data. The sequences were then clustered into operational taxonomic units (OUTs) at 97% similarity. The results were further analysed on the online platform of Majorbio Cloud Platform including plotting figures.
